# Do native and introduced cover crops differ in their ability to suppress weeds and reduce seedbanks? A Case study in a Table Grape Vineyard

**DOI:** 10.1101/2023.06.20.545823

**Authors:** Margaret R. Fernando, Lauren Hale, Anil Shrestha

## Abstract

Cover cropping is considered a valuable tool for integrated weed management. However, weed suppression by cover crops can be inconsistent. We hypothesized that a native cover crop species may have greater capacity to suppress weeds than an introduced species owing to adaptive advantages. A study was conducted from 2020 to 2022 in a newly-established Autumn King tablegrape vineyard in Parlier, CA. Two cover crop treatments, a native plant species, phacelia (*Phacelia tanacetifolia*), or an introduced species, rye (*Secale cereale* L. ‘Merced’), were compared to a no cover crop, herbicide-managed control for impacts on weed control. Cover crops were sown in 2019 in the center 1.8 m of the 3.6 m space between the grapevine rows. The experimental design was a randomized complete block with four replications. Percent cover, weed density, and weed biomass in different seasons were assessed in the inter-row spaces and the vine rows. Soil samples were collected in 2021 from the grape row and interrow spaces to assess the weed seedbank. After the cover crops were terminated, percent weed cover was lower in the interrow of the phacelia compared to the no cover crop plots at two sampling times. When cover crops were actively growing (winter/spring), phacelia plots had a 7-fold higher cover crop to weed biomass ratio compared to that of the rye plots, thus indicating greater weed suppression. However, weed seedling emergence from the seedbank samples of the cover crop plots were 2- to 4-fold greater (in the phacelia and rye, respectively) than from the no cover crop plots. In summary, phacelia suppressed weeds more than rye; however, they both resulted in a bigger weed seedbank size compared to the herbicide-managed no cover crop plots, so additional management practices will be needed for long term weed management with cover crops in vineyards.

## Introduction

Weed management generally represents a significant portion of farm budgets in agricultural cropping systems (Gianessi and Reigner 2007). Although growers commonly tend to rely on herbicides as a convenient and inexpensive tool for weed control, these chemicals can pollute surface water and groundwater in addition to altering soil characteristics (Aktar et al. 2009). A study reported that 41% of the 1204 wells sampled across several regions of the US contained pesticides (Bexfield et al. 2021). Another study reported the negative effect of herbicides on soil quality caused by shifts in soil microbial communities and declines in mycorrhizal fungal growth (Chakravarty and Sidhu, 1987; Druille et al. 2013). Such effects can have negative implications to crop productivity as mycorrhizal fungi form mutualistic, symbiotic relationships with plant roots and are considered large components of ecosystem functioning and plant health (van der Heijden et al. 1998). Overuse of herbicides has also influenced the evolutionary processes in weeds leading to herbicide resistance mechanisms in weeds (Powles and Yu 2010). Current trends in weed science demonstrate a search for a holistic, sustainable approach, which focuses on integrating different tools for weed management instead of the sole reliance on herbicides (Shrestha et al. 2004). Strategies, such as the use of cover crops, may be one such approach as they may enhance farm sustainability by providing resource-efficient and cost-effective solutions to address environmental concerns associated with common weed control practices as well as improve soil health. Many studies have demonstrated soil health benefits of cover crops including higher C and N pools, microbial biomass, and aggregate stability (Belmote et al., 2018; Stine et al., 2002), and some studies have also indicated weed suppression effects of cover crops (Hoffman et al., 1996; Caamal-Maldonado et al., 2001). Cover crop adoption rate in the United States, however, is much lower in California than in the midwest and northeast states (LaRose and Myers 2019). This could be due to the concern of water use by cover crops during the winter months hence causing moisture stress to the cash crop during the growing season (Mitchell, Shrestha, and Irmak 2015). Although in the case of vineyards, in some areas where rainfall is more prevalent, cover crops may be planted under grapevines to reduce excess vine vigour by allowing competition for water (Heuvel, 2017). However, this is generally not the case in sandy soils in the semi-arid region of the Central Valley of California. Due to recent shifts in acreage from annual to perennial systems in the Central Valley, there has been an increase in the minimum irrigation needs (Mall et al. 2019; Perdue and Hamer 2019). Because of the predicted increase in frequency of drought (IPCC 2014), the water requirements of cover crops have become an obstacle when promoting the inclusion of cover crops in cropping systems; however, if cover crops cause the suppression of weed populations, water conservation could potentially be promoted.

Most of the species chosen as cover crops have focused on several introduced species that include grasses and legumes. However, the use of native species as cover crops have not been adequately explored. We hypothesized that a native species of California such as phacelia (*Phacelia tanacetifolia*), because of its adaptation to local conditions, would have better weed suppressive abilities than an introduced species such as rye (*Secale cereale* L.). Therefore, a study was conducted to assess the differences in weed suppression capacity between phacelia and rye when used as cover crops in a table grape vineyard in comparison to no cover crop. Specific objectives of the study were to: a) evaluate the differences between the native (phacelia) and introduced (rye) cover crops and no cover crop (control) treatments on weed densities in the interrow spaces of the vineyard, b) compare the difference between treatments on weed populations and biomass, and c) to assess the weed seedbank size in the three treatments.

## Materials and methods

The study was conducted from winter 2019 to summer 2022 to evaluate the effects of a native and an introduced cover crop in a table grape vineyard in Parlier, CA (36.5960049 W, - 119.5119173N). The study site is located in the southern part of the Central Valley of California where the climate is Mediterranean type with very hot and dry summers and cool and damp winters. The average temperature is highest (approximately 37°C) in July and lowest (approximately 13° C) in December. The average annual rainfall is approximately 263 mm with March being the wettest month (Weather US, 2023). The soil at the site was a Hanford sandy loam (Coarse-loamy, mixed, superactive, nonacid, thermic Typic Xerorthents). Cover crop seeds were broadcast by hand and raked into the soil in December 2019. The following seasons (2020 and 2021) the cover crops were self-maintained from their own seed production and shed, occurred after the end of their respective life cycles, thus, plots were not reseeded. Autumn King grapevines on Freedom variety rootstock were planted in May 2020. Before planting the cover crops and grapevines, the plots were disked, but not tilled thereafter. A modified T cable trellis system was used to support the grapevines, and sleeves were applied to the young plants for protection from pests. A cane pruning strategy adapted from El-Kereamy and Kurtural (2022) for use on a T cable trellis was used in the study. The treatments included Lacy phacelia as native cover crop species and rye cv Merced as introduced cover crop species; a no cover crop control treatment was also included. Each treatment plot was 22 m long and 11 m wide and included four rows of grapevines and three rows of cover crops in the interrow spaces of the vine rows. The experimental design was a randomized complete block, with four repetitions, where three rows of grapevines (running east to west) were a block, and the block was subdivided into three plots in which the treatments were randomized.

Cover crops were planted in the centre 1.8 m of the 3.6 m total space between the grapevine rows. Seeds were broadcast at a rate of 8.9 and 6.7 kg ha^-1^ for phacelia and rye (Kamprath Seed Co., Manteca, CA), respectively. The grapevine rows were irrigated via metered drip irrigation, and cover crop rows were irrigated with metered, micro-sprinkler irrigation. The cover crops had to be irrigated for establishment because of lack of rain (63% and 71% of average precipitation in Nov. 2019-March 2020 and Nov. 2020-March 2021, respectively). The area under the grapevines in all the plots and inter-row spaces of the control plots were sprayed with the broad-spectrum, postemergence herbicide glufosinate-ammonium (Lifeline, Glufosinate Ammonium 280 SI, UPL NA Inc., King of Prussia, PA), at 3.5 l ha^-1^ in April 2020, May2020, and February 2021; the area under vine rows was sprayed again in August 2020. Additional applications of glufosinate-ammonium were made during July 2020 and November 2020 to the inter-row spaces and the area under the grapevines of all treatments. Herbicide applications under the grapevine rows and interrow spaces were made based on the standard practice in table grape vineyards in the Central Valley.

Cover crops were mown on June 16, May 17, and March 11 in 2020, 2021, and 2022, respectively. These mowing dates were based on the height of the cover crops that varied between the years due to seasonal differences in weather conditions. The mowing took place after the cover crops had seeded. The mown residue was left in the plots to serve as a mulch. A pesticide spray program including a rotation of several fungicides and insecticides were applied to vine foliage approximately every 20 days and gopher traps and an owl box were set up to control gopher populations.

To assess the capacity of cover crops to control weeds both in interrow spaces and beneath vines, weed pressure estimates were collected independently in interrow spaces and in grapevine rows in all treatment and control plots. Although the area under the grapevine was treated similarly in all the plots, weed populations were assessed to verify if encroachment of the cover crop roots in this area would alter soil moisture dynamics and hence weed emergence and coverage and also if seeds from the cover crops and weeds would encroach this area by various dispersal mechanisms. Weed populations were compared between the treatments using visual estimates of total percent of ground cover by weeds and by weed species to estimate the density. The soil weed seedbank size was measured by taking soil cores, and weed suppression by the cover crop was also estimated by taking samples of weeds and cover crop biomass and calculating covercrop:weed biomass ratio. For percent ground cover by weeds and weed species density, a 0.5 m x 0.5 m polyvinyl chloride (PVC) quadrat was randomly tossed eight times in each plot (four times each within the grapevine row and in the interrow spaces). The percent ground cover by weeds was estimated for each toss, and individual weeds within the quadrat was counted and identified by species. This evaluation was performed several times during the year in different seasons. All the weed samplings to measure percent ground cover by weeds in the interrow spaces were taken after the cover crops had been mowed, and the residues had been left to act as a mulch.

The soil weed seedbank was assessed once during the experiment in winter 2021 using the greenhouse germination method described by Shrestha et al. (2015). A 7-cm diameter soil auger was used to collect soil samples from 0-6 cm and 6-20 cm depths at four random spots in the third row under the grapevines and four random spots in the cover cropped interrow area. The four samples in the vine row were bulked together and the four samples from the interrow area were bulked together for a total of 48 samples (12 plots x 2 depths x 2 locations). Samples were stored in a greenhouse set at a constant temperature of 25° C and spread in a thin (0.5 cm) layer on 30 cm diameter plastic trays. The soil was watered regularly to stimulate seed germination, and each week the emerged weed seedlings were counted by species, recorded, and removed; the seedbank was monitored for approximately six months. To prevent soil compaction, soil was stirred once a month. To assess the suppressive effect of cover crop on weeds during the actively growing season of the cover crops, samples of weed and cover crop biomass were collected during spring 2022. A 0.5 m x 0.5 m PVC quadrat was tossed randomly twice per plot in each replication. The plants growing inside the area of the quadrat were clipped at the soil surface, collected, bagged, transported to the lab, and separated into weeds or cover crops. The biomass of the separated species was then dried in a forced-air oven at 40°C for a two weeks and dry weights were recorded.

Data were analysed using ANOVA or ANOVA on Ranks using Sigma Plot (version 14.0). Data collected in the grapevine rows and the interrow spaces were analysed separately. Assumptions of ANOVA (normality and homogeneity of variance) were checked prior to the analysis. If the assumptions were not met, appropriate transformations were performed. When the transformed data still did not pass the assumptions, data were analysed using a nonparametric test (Kruskal-Wallis One Way Analysis of Variance on Ranks). Because this test measures significant differences between median values of treatments, medians along with means were presented in tables when this test was used. When the ANOVA indicated significant differences at a 0.05 alpha level, Holm Sidak method, Dunn’s method, or Tukey’s Test was used for multiple comparison between the treatments. The weed species information from the samplings was organized in a species table, and it was plotted using principal component analysis (PCA). R (version 4.2.1) vegan and ggplot2 packages were used for composition of weed family heatmap across time as well as for PCA analysis of weed species in the grapevine row and interrow spaces.

## Results

### 1. Percent Ground Cover by Weeds

Percent ground cover by the weeds in the interrow spaces was higher in the control without cover crop than in phacelia in the first year (Table 1), with the control having 51% more weed cover compared to phacelia plots (P=0.038). However, no significant differences were observed between rye cover crop and control plots. Phacelia plots also had lower total weed density in the interrow spaces compared to the other treatments (*P*=0.031) (Table 2). Although the percent ground cover by weeds in the grapevine rows was similar between the different treatments (*P*=0.294), there were differences in the dominant species. Common knotweed (*Polygonum aviculare*) densities were higher in the grapevine row as well as in the interrow spaces (P=0.015) of rye and control plots compared to the densities in the phacelia plots. The average common knotweed densities within the grapevine row were 6±2, 15±3, 17±1 plants m^-2^ for the phacelia, rye, and no cover crop plots, respectively; the average densities in the interrow spaces were 5±1, 23±1, 15±1 plants m^-2^, respectively for these treatments. This weed species was the most prominent in all the seasonal assessments of species density (Figure 1). Horseweed (*Erigeron canadensis*) was the second most common species found in the weed samplings, however, there was no significant difference in its densities between the three treatments. In spring and summer 2021, percent ground cover of the weeds was generally lower than in fall 2020, and there were no differences between the treatments (P>0.05). By spring 2022, percent ground cover of the weeds in the interrow spaces was higher (32%) in the control compared to the cover crop treatments (6%) (P<0.001). However, there were no differences between treatments in the percent of weed cover in summer 2022 in both vine row and interrow spaces (P=0.658, P=0.134, respectively). Regarding the weed species density, there seemed to be more similarity between treatments within grapevine rows compared to the interrow areas (Figure 2).

**Figure 1.**
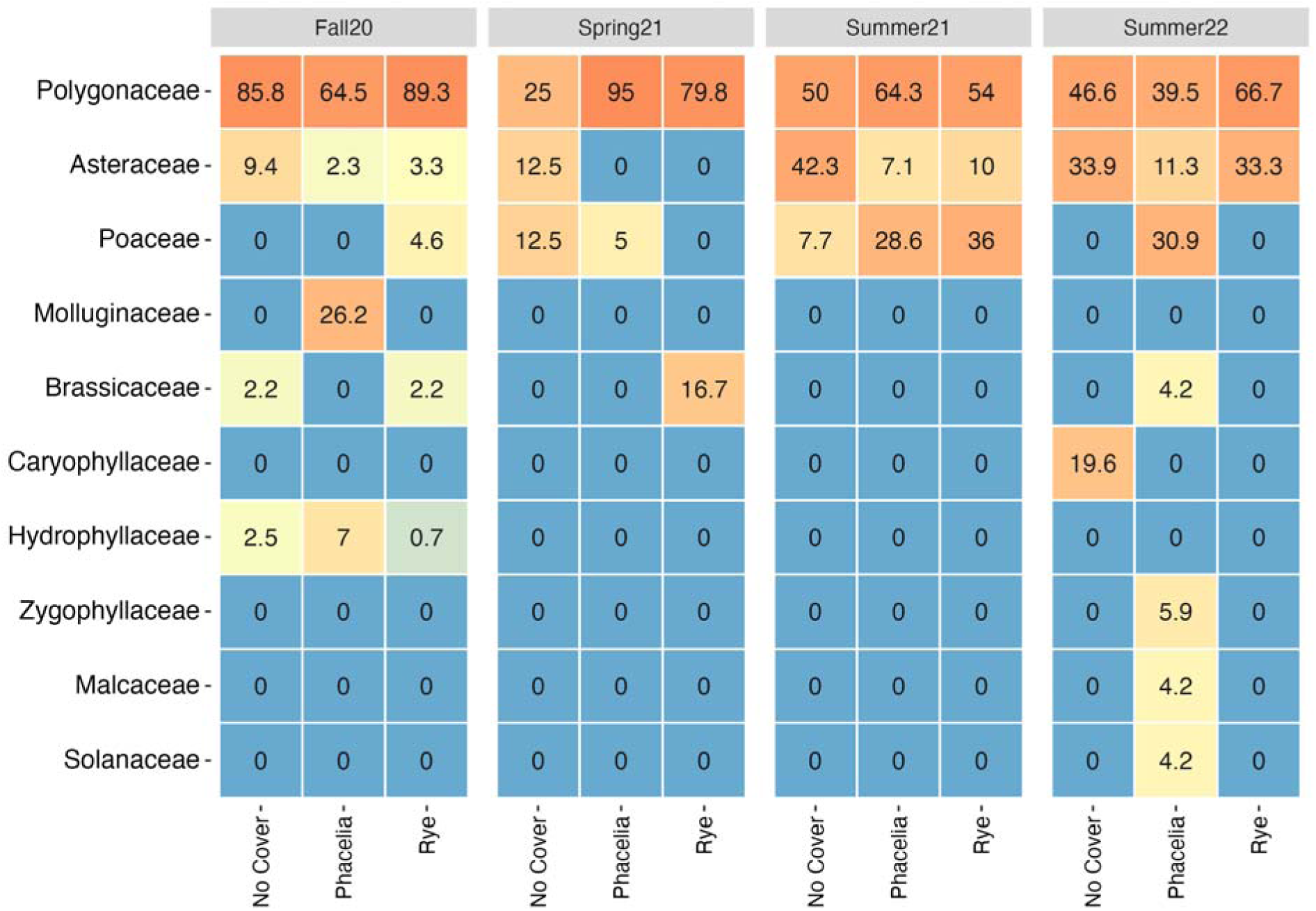
**Seasonal distribution of weed families found in native (phacelia) species cover crop, introduced (rye) species cover crop, and no cover crop treatment plots in both grapevine rows and in the interrow spaces. Data are sorted based on average percent abundance of weed families in the three treatments during each seasonal sampling.**

**Figure 2.**
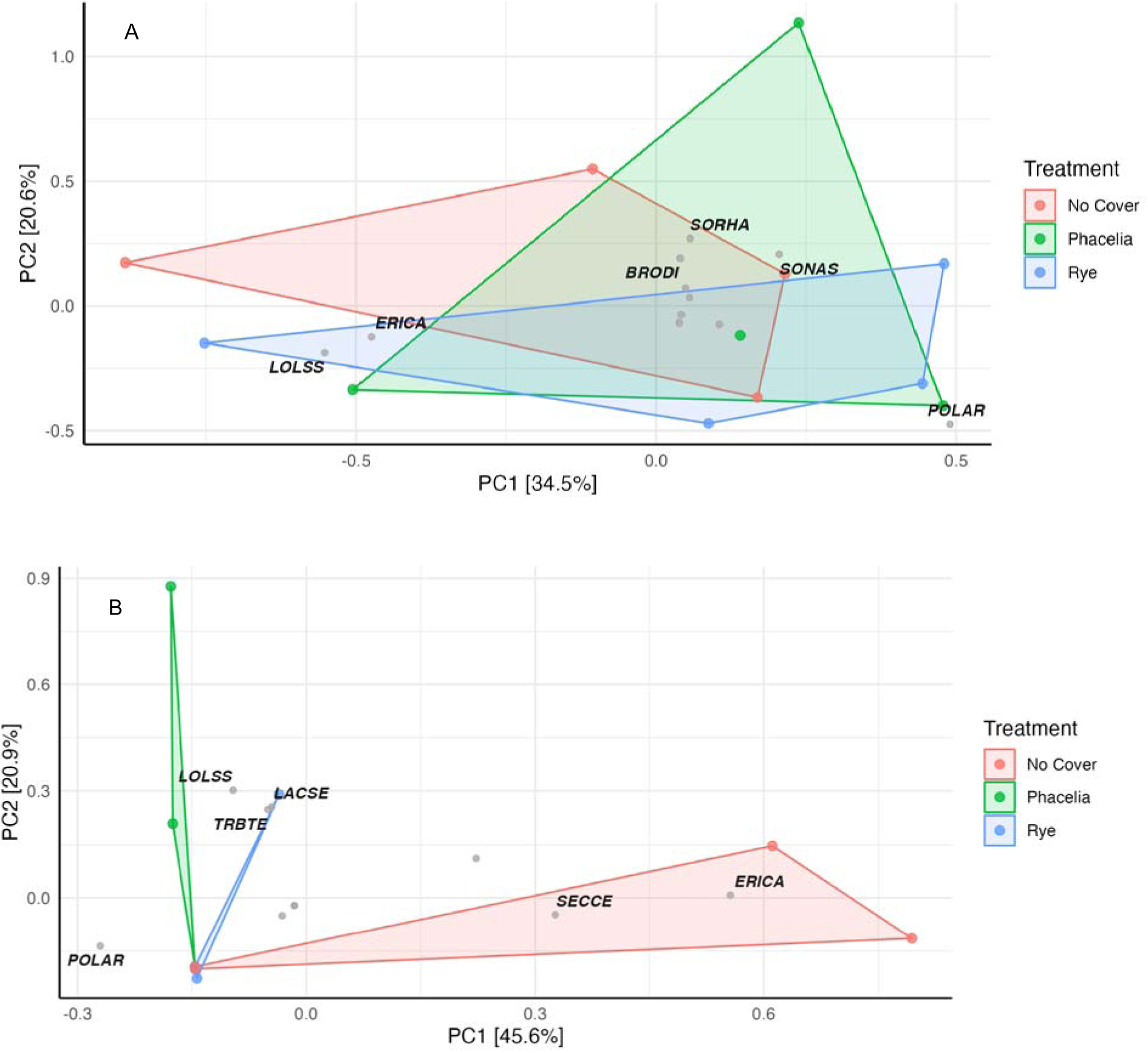
**Principal Component Analysis (PCA) for weed species in the grapevine rows (A) compared to the interrow spaces (B). Bayer codes for weed species: common knotweed = POLAR, ryegrass=LOLSS, horseweed= ERICA, ripgut brome=BRODI, Johnsongrass= SORHA, annual sowthistle= SONAS, rye=SECCE, puncturevine= TRBTE, prickly lettuce=LACSE.**

**Table 1.** **Percent weed cover determined during three establishment years of an Autumn King table grape vineyard. Values represent means with standard error along with median values for percent weed cover collected fall 2020 through summer 2020 from 4 replicate plots.**

| Location | Treatment | Fall 2020 |  | Spring 2021 |  | Summer 2021 |  | Spring 2022 |  | Summer 2022 |  |
| --- | --- | --- | --- | --- | --- | --- | --- | --- | --- | --- | --- |
|  |  | Mean | Median | Mean | Median | Mean | Median | Mean | Median | Mean | Median |
|  |  | % | % | % | % | % | % | % | % | % | % |
| Vine rows | Phacelia | 6 ± 2 | 3 | 0 ± 0 | 0 | 4 ± 2 | 4 | n.a. | n.a. | 3 ± 1 | 4 |
|  | Rye | 4 ± 1 | 3 | 2 ± 2 | 0 | 16 ± 7 | 1 | n.a. | n.a. | 2 ± 1 | 1 |
|  | No Cover | 3 ± 2 | 1 | 0 ± 0 | 0 | 1 ± 0 | 10 | n.a. | n.a. | 2 ± 1 | 10 |
| Interrow spaces | Phacelia | 22 ± 8 | 7<sup>a</sup> | 2 ± 1 | 1 | 12 ± 6 | 4 | 6 ± 1 | 5<sup>a</sup> | 11 ± 3 | 1 |
|  | Rye | 35 ± 9 | 23<sup>ab</sup> | 3 ± 1 | 1 | 8 ± 3 | 0 | 6 ± 2 | 5<sup>a</sup> | 2 ± 1 | 0 |
|  | No Cover | 43 ± 8 | 38<sup>b</sup> | 2 ± 1 | 0 | 2 ± 1 | 0 | 32 ± 5 | 28<sup>b</sup> | 4 ± 1 | 0 |

\* Treatment medians in a column within location and season followed by different lower-case letters are significantly different from each other according to Tukey test at a 0.05 level of significance. n.a. = not available

**Table 2.** **Total weed species density determined during three establishment years of an Autumn King table grape vineyard in Parlier, CA. Values represent means with standard error for total weed species density collected fall 2020 through summer 2020.**

| Location | Treatment | Fall 2020<br>Plants m<sup>-2</sup> | Spring 2021<br>Plants m<sup>-2</sup> | Summer 2021<br>Plants m<sup>-2</sup> | Summer 2022<br>Plants m<sup>-2</sup> |
| --- | --- | --- | --- | --- | --- |
| Vine rows | Phacelia | 21 ± 14 | n.a. | 2 ± 0 | 4 ± 1 |
|  | Rye | 17 ± 2 | n.a. | 2 ± 1 | 3 ± 1 |
|  | No Cover | 23 ± 2 | n.a. | 4 ± 2 | 6 ± 2 |
| Interrow<br>spaces | Phacelia | 7 ± 1<sup>a</sup> | 4 ± 1 | 4 ± 1 | 4 ± 1 |
|  | Rye | 14 ± 1<sup>b</sup> | 3 ± 1 | 1 ± 1 | 5 ± 2 |
|  | No Cover | 15 ± 1<sup>b</sup> | 1 ± 0 | 1 ± 1 | 6 ± 2 |

\* Treatment means in a column within location and season followed by different lower-case letters are significantly different from each other according to Holm-Sidak method at a 0.05 level of significance.

### 2. Weed Seedbanks

The total number of weed seedlings that emerged from the soil seedbank samples collected from the interrow spaces were higher in the cover crop plots than from the control plots (P=0.001). The average number of seedlings in the 0-5cm layer were 870, 1740, and 414 m^-2^ for phacelia, rye, and control plots, respectively (Table 3). The number of seedlings emerged from the 5-20 cm layer samples followed the same trend (Table 4) with both cover crop treatments in the interrow spaces showing higher total number of seedlings the control without cover. The two cover crop treatments (phacelia and rye) differed from each other in the individual species density. Within the interrow spaces of the 0-5 cm layer (Table 3), phacelia had more common knotweed seedlings (P=0.004), while the rye cover crop had more common chickweed (*Stellaria media*) seedlings (P=0.018). However, there were no differences in the number of emerged weed seedlings between the three treatments in the soil samples collected from the grapevine rows (P=0.609). Weed seedling emergence in all the treatment samples was greater from the shallow soil depth (0-5 cm) than from the deeper soil depth (5-20 cm) in the interrow spaces with an average of 61 and 44 plants m^-2^, respectively (P<0.001). However, such differences did not occur between the two soil depths in the samples from the grapevine rows (P=0.092).

**Table 3.** **Average number (±SE) of seedling emergence of the dominant species (common knotweed and common chickweed) from soil samples taken from the 0-5 cm depth, during the third year (Winter 2021) of establishment in an Autumn King table grape vineyard in Parlier, CA.**

| Location | Treatment | Total<br>Plants m<sup>-2</sup> | Common knotweed<br>(*Polygonum*<br>*aviculare*)<br>Plants m<sup>-2</sup> | Common<br>chickweed<br>(*Stellaria media*)<br>Plants m<sup>-2</sup> |
| --- | --- | --- | --- | --- |
| Vine row | Phacelia | 205 ± 36 | 50 ± 32 | 127 ± 42 |
| | Rye | 245 ± 36 | 25 ± 21 | 212 ± 85 |
| | No Cover | 191 ± 41 | 7 ± 4 | 156 ± 29 |
| Interrow<br>spaces | Phacelia | 870 ± 116 <sup>a</sup> | 177 ± 35 <sup>a</sup> | 594 ± 85 <sup>a</sup> |
| | Rye | 1740 ± 435 <sup>a</sup> | 24 ± 12 <sup>b</sup> | 1556 ± 410 <sup>b</sup> |
| | No Cover | 414 ± 134 <sup>b</sup> | 50 ± 21 <sup>b</sup> | 255 ± 127 <sup>a</sup> |

\*Means within a column for each location (vine row or interrow) followed by different lower-case letters are significantly different from each other according to Holm-Sidak method at a 0.05 level of significance. Mean separation for interrow values were based on square root-transformed data, but non-transformed means are presented.

**Table 4.** **Average number (±SE) of seedling emergence of the dominant species (common knotweed and common chickweed) from soil samples taken from the 5-20 cm depth, during the third year (Winter 2021) of establishment in an Autumn King table grape vineyard in Parlier, CA.**

| Location | Treatment | Total Weed<br>Density<br>Plants m<sup>-2</sup> | Common knotweed<br>(*Polygonum*<br>*aviculare*)<br>Plants m<sup>-2</sup> | Common chickweed<br>(*Stellaria media*)<br>Plants m<sup>-2</sup> |  |
| --- | --- | --- | --- | --- | --- |
|  |  | Mean | Mean | Mean | Median |
| Vine rows | Phacelia | 118 ± 37 | 7 ± 7 | 57 ± 23 | n.a. |
| | Rye | 158 ± 37 | 35 ± 17 | 90 ± 50 | n.a. |
| | No Cover | 59 ± 43 | 5 ± 4 | 39 ± 35 | n.a. |
| Interrow spaces | Phacelia | 294 ± 35 <sup>a</sup> | 35 ± 26 | 223 ± 21 | 205 <sup>ab</sup> |
| | Rye | 474 ± 157 <sup>a</sup> | 25 ± 9 | 421 ± 144 | 417 <sup>b</sup> |
| | No Cover | 99 ± 30 <sup>b</sup> | 18 ± 11 | 57 ± 39 | 28 <sup>a</sup> |

\*Means within a column for each location (vine row or interrow) followed by different lower-case letters are significantly different from each other according to Holm-Sidak method at a .05 level of significance. Mean separation for interrow values were based on square root-transformed data, but non-transformed means are presented. n.a. = not available.

### 3. Weed Biomass and Ratio of Cover Crop to Weed Biomass

The weed biomass in the interrow spaces was greater in rye plots than in phacelia and control plots (P=0.008). The average weed biomass in phacelia and control plots was approximately 86% and 84% lower than that in the rye cover crop plots, respectively (Table 5). The ratio of cover crop biomass to weed biomass was greater in phacelia than in the rye, with an average of approximately 33 and 5 g m^-2^, respectively (P=0.005). The weed biomass in the grapevine rows was similar between the three treatments (P=0.294).

**Table 5.** **Weed biomass and cover crop:weed biomass ratio during Spring 2022 in an Autumn King table grape vineyard in Parlier, CA.**

|  |  | Weed Biomass<br>g m<sup>-2</sup> |  | Cover:Weed Biomass |  |
| --- | --- | --- | --- | --- | --- |
|  |  | Mean | Median | Mean | Median |
| Vine rows | Phacelia | 32.9 ± 19.2 | 8.2 | n.a. | n.a. |
|  | Rye | 22.5 ± 9.5 | 13.6 | n.a. | n.a. |
|  | No Cover | 62.6 ± 22.7 | 43.2 | n.a. | n.a. |
| Interrow spaces | Phacelia | 13.0 ± 4.9 | 6.8 <sup>a</sup> | 33.3 ± 9.5 | 27.7 <sup>a</sup> |
|  | Rye | 93.0 ± 29.5 | 56.6 <sup>b</sup> | 5.3 ± 1.7 | 4.0 <sup>b</sup> |
|  | No Cover | 15.2 ± 6.0 | 8.8 <sup>a</sup> | n.a. | n.a. |

\*Means with standard error and medians for weed and cover crop biomass in the grapevine rows and interrow spaces in a column within location (vine rows or interrow spaces) followed by different lower-case letters are significantly different from each other according to Tukey test at a 0.05 level of significance. n.a. = not available.

## Discussion

In this study, postemergence herbicides were applied at similar rates in Spring in the grapevine rows of all treatments, and there were no cover crops in the grapevine rows. Thus, the weed management system was similar in all the treatments in the grapevine rows. Therefore, this explains why weed densities, percent weed cover, weed biomass, and seedbank estimates were similar between the treatments in the grapevine rows. The percent ground cover of weeds and weed densities in the interrow spaces of phacelia plots were lower than in the no cover and in the rye plots in Fall 2020, and thus, proving to be the most effective among the three treatments for weed suppression. In particular, the densities of common knotweed, the most abundant weed species, were lowest in phacelia plots indicating its higher capacity of suppressing this species than rye. Teasdale and Mohler (2000) also reported differences between cover crop species on weed suppression. However, the exact reason for such differences is not known. Numerous studies have reported suppression in weed emergence by a rye cover crop (Teasdale and Mohler, 1993; Mirsky et al. 2013; Grint et al. 2022). This could partly be due to the allelopathic properties of rye as described by Barnes and Putnam (1987) and reduced soil temperature and reduced light and moisture availability under rye residue (Mirsky et al., 2013; Teasdale and Mohler, 1993). In comparison, very few studies have been conducted on the allelopathic properties of phacelia. One study reported suppression of some Asteraceae weed species by aqueous extracts of phacelia (Puig et al., 2021). Another study in Germany (Brust et al. 2014) reported reduction in weed densities by up to 77% by phacelia, and they hypothesized that this was because of the same factors described above for rye. Similarly, phacelia residues were reported to reduce weed emergence when used as a cover crop before cabbage planting (Franczuk et al. 2009). In our study, seasonal measurements demonstrated the success of the native cover crop species in suppressing weed populations; moreover, the introduced cover crop species, rye, did not show any obvious suppressive effects but was able to maintain similar weed density levels without herbicides as the control plots where herbicides were applied. This may be attributed to the greater biomass produced by phacelia compared to rye (data not shown).

Unlike in the Fall, no differences between the treatments in percent weed cover was observed in evaluations taken in Spring 2021 and Summer 2021, both in the grapevine rows and in the cover crop rows. This could be due to the especially dry winter of that year, as weed populations were, in general, relatively low in these measurements. However, in Spring 2022, the percent weed cover in the interrow was higher in the control treatment than in the cover crop plots, thus, during this season, the cover crops weed suppressive capacity was similar, and more effective than that of the control plots.

In summer 2022, differences in percent ground cover in the interrow spaces between the treatments disappeared. This probably was because most of the weeds had emerged in spring and covered the ground. While some studies have reported reduction in weed populations with cover crops (Nichols et al., 2020; Matloob and Chauhan, 2021), there are studies that have reported no effect of cover crops on weed densities (Reddy, 2001; Swanton et al., 1999). Thus, the effects of cover crops on weed suppression have been inconsistent and dependent on several physical, biological, and environmental factors. In our study, the effect of cover crop treatment on percent cover in the interrow spaces varied from season to season. Longer term experiments comparing the three treatments may provide a better picture if the trend in seasonal differences holds true and if cover cropped plots suppress weeds consistently in the interrow row spaces compared to the uncropped control plots.

Lowering the soil weed seedbank is an important integrated weed management strategy because the persistence of weeds in the field is dependent on the size of the soil seedbank (Schwartz-Lazaro and Copes 2019). The cover crop treatments did not affect the weed seedbank size of the grapevine rows because the amount of weed seedling emergence from the soil seedbank samples was similar between them; however, this was expected because additional herbicide treatments were applied in the grapevine rows in all the treatment plots. In the interrow spaces, the cover crop treatment plots resulted in an increase in the weed seedbank size because there were more weed seedling emergence from the samples collected from the cover crops plots compared to the no cover crop treatment plots. Shrestha et al. (2015), also reported a similar phenomenon in terms of a larger weed seedbank in plots with cover crops compared to no cover crops. This could be because, while broad spectrum postemergence herbicides kill many weeds and reduce weed seed production, several weed species can continue growing in the understory of the cover crops and produce seeds successfully and add to the seedbank. Although Nichols et al. (2020) concluded that long-term use of cover crops in a maize *Zea mays*) and soybean (*Glycine max*) rotation had the potential to reduce the size of weed seedbanks compared to winter fallow, their conclusion may have been different if herbicides were applied on the weeds during the winter fallow. The success of cover crops in reducing weed seedbank size in herbicide-free systems has also been reported by Schmidt et al. (2019). A review by Sias et al. (2021) on the effects of cover crop on weed seedbanks was inconclusive and suggested that future research should also focus on factors such as dormancy status, predation levels, weed species composition, and germination to get a better idea of weed seedbank dynamics. It can be inferred from our study that, in the long-term, use of cover crops alone without supplemental weed control methods may increase the size of the weed seedbank in the interrow spaces of the vineyards. This inference is also indirectly supported by the conclusions of Baraibar et al. (2017) who emphasized the importance of using competitive cover crops to prevent or reduce weed seed rain from escapes that could add to the size of the soil seedbank adding to future weed problems.

There were differences between the weed species compositions in the seedbank samples from phacelia and rye. Phacelia showed had higher number of common knotweed seedlings, while rye had higher number of common chickweed seedlings. The reason for this difference is not certain but phacelia seemed to favour common knotweed which is a species with a prostrate growth habit while rye favoured the more erect growing common chickweed species which tended to climb on rye plants. Difference in light attenuation patterns through the canopy of the two cover crop species could have caused this phenomenon. However, light measurements were not taken in this study. This suggests that the suppression of a particular weed species may depend on the cover crop species, in agreement with past research from Creamer et al. (1996). Brennan and Smith (2017) also reported that mustard cover crops reduced seed production of burning nettle (*Urtica urens*) more than oats (*Avena sativa*) or legume/oats mixtures. The assessments of the weed seedbank also showed higher seedling emergence in soil samples collected from shallow depth (0-5 cm) compared to the deeper depth (5-20 cm). This can be attributed to the fact that weed seeds were not incorporated into the soil as tillage was only done in the interrow spaces during the first year of the project. The subsequent years were basically no-till systems. Swanton et al. (2000) reported that, in sandy soils, as in the case of our study, 90% of the weed seeds were confined to the 0-5 cm of the soil depth.

In this study, the weed assessments of percent cover and density were not taken during the active growing season of the cover crop to reduce confusion of equating plant cover with weed cover. While it is interesting to observe how the cover crop residues affected weed populations, it was also important to document how the actively growing cover crops affected weed populations. Because plant competitiveness can be explained by biomass traits (Gaudet and Keddy, 1988), in our study, above ground biomass samples of both the weeds and the cover crops were taken before the cover crops were mowed. We found that in early spring 2022, phacelia and control treatments had lower weed biomass compared to rye. It was also observed that the cover crop biomass to weed biomass ratio was higher in phacelia with respect to rye. This implies that the native species, phacelia, was more successful in competing against weeds compared to the introduced species, rye. Teasdale and Mohler (1993) and Teasdale (1993) reported that biomass produced by cover crops affected the germination of weed seeds by creating shaded areas, so it could be possible that phacelia created more shaded area compared to rye, causing a reduction in weed populations. Therefore, in this study, phacelia was a better choice than rye in terms of weed suppression. However, it is not known if the same holds true in other cropping systems and other locations. Regardless, this study also showed that cover crops alone may not be sufficient to suppress weeds and reduce weed seedbank sizes in interrow spaces of vineyards in the short term. A recent study has also suggested that while cover crops have value in weed suppression, they do not have a consistent effect compared to repeated herbicide applications (Haring and Hanson, 2022). Supplementary weed control measures may need to be adopted in the interrow spaces to lower the weed seedbank and the environmental and economic benefits of using cover crops compared to herbicides may need to be evaluated for sustainability assessments.

## Practical Implications

This study showed that, when planted as cover crops in the interrow spaces of grape vineyard, the native species phacelia, in general, had more favourable outcomes on weed management than the introduced rye or the herbicide-managed control plots in a Central Valley tablegrape vineyard. However, the interrow spaces of both cover crop treatments had a bigger weed soil seedbank size as, estimated by the seedling emergence method, compared to the control plots. This could be an indication that weed populations in the cover crop treatments may increase over time without supplemental weed management interventions. Although the effects of cover crops on weed populations may shift from season to season, cover crops, especially the native species, seemed as effective as the no cover crop treatment. This is an indication that native species may have better potential than introduced species as cover crops for weed suppression in agricultural cropping systems. If cover crops keep reducing weed biomass and weed seed production, weed densities may be lowered in the long-term making them a very effective tool in integrated weed management in table grape vineyards in the San Joaquin Valley of California.

## Acknowledgements

Funding for this project was provided by the USDA-ARS, California Department of Food and Agriculture Healthy Soils Program, and UC SAREP. We would like to thank Natalie Scott at the USDA-ARS for helping process plant samples and for helping set up the weed seedbank experiment.

